## Supplementary Fig 1 for "Adeno-associated virus capsid assembly is divergent and stochastic"

a

3.7 MDa

| z | $m/z_c$ | σ | $I_c$ |
|---|---|---|---|
| 147 | 25,171 | 9.1 | 0.3 |
| 148 | 25,001 | 9.0 | 0.6 |
| 149 | 24,833 | 8.9 | 0.8 |
| 150 | 24,668 | 8.8 | 1.0 |
| 151 | 24,504 | 8.7 | 0.8 |
| 152 | 24,343 | 8.6 | 0.6 |
| 153 | 24,184 | 8.5 | 0.3 |

$$I(m/z) = \frac{I_c}{\sigma\sqrt{2\pi}} e^{-\frac{1}{2}\left(\frac{m/z - m/z_c}{\sigma}\right)^2}$$

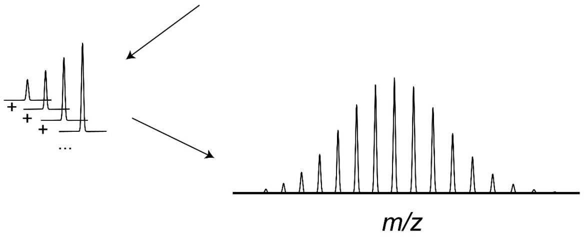


b

8:8:84 (VP1%:VP2%:VP3%)

| VP1:VP2:VP3 | Mass | p | z_average |
|---|---|---|---|
| 0:0:60 | 3,568,500 | $2.86x10^{-5}$ | 147 |
| 0:1:59 | 3,575,118 | $1.64x10^{-4}$ | 147 |
| 1:0:59 | 3,590,269 | $1.64x10^{-4}$ | 147 |
| 1:1:58 | 3,596,887 | $9.19x10^{-4}$ | 148 |
| 0:2:58 | 3,581,736 | $4.60x10^{-4}$ | 147 |
| 2:0:58 | 3,625,274 | $6.89x10^{-3}$ | 148 |

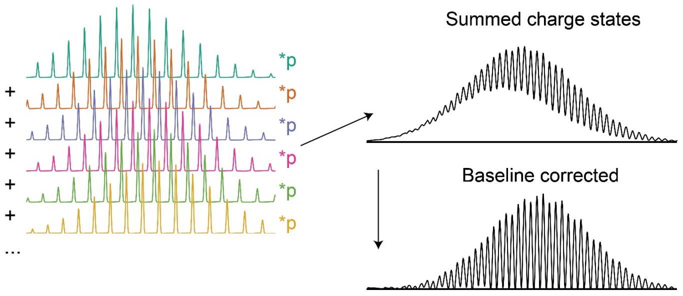


c

| VP1%:VP2%:VP3% | Average Mass |
|---|---|
| 0:0:100 | 3,568,500 |
| 0:1:99 | 3,572,471 |
| 1:0:99 | 3,581,561 |
| 1:1:98 | 3,585,532 |
| 0:2:98 | 3,576,442 |
| 2:0:98 | 3,594,623 |

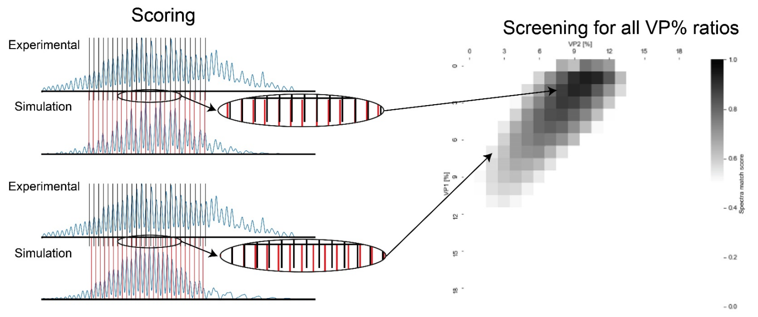


***Supplemental Figure 1 - Overview of simulation and scoring workflows: a)*** *Illustration of procedure for simulation of individual charge state distributions. For each charge state we calculated the corresponding m/z-position, resolution dependent peak width, and normally distributed intensity values. Using the displayed formula, each peak was calculated separately and combined to the final charge state distribution.* ***b)*** *For the Simulations of whole AAV populations we first calculated the mass distribution for all 1891 possible VP stoichiometries as shown in Figure 4a. With the calculated average charge and a fixed charge state width we calculated each charge state distribution as described in* ***a)*** *and multiply them by their probability before combining them into one final mass spectrum. For better comparability with transient averaged mass spectra we performed a baseline correction step.* ***c)*** *For the systematic comparison between simulations and experimental data we changed the bulk VP% expression levels by increments of 1%. From the average mass we adjusted the charging so the simulated charge state distribution populates the same m/z regions as the experimentally recorded spectrum. The peak positions are then compared and scored as described in the method section. The top spectra illustrated a good match between simulation and experimental data whereas the bottom spectra is an example of a bad match with diverging peaks positions.*
